# Single-dimensional human brain signals for two-dimensional economic choice options

**DOI:** 10.1101/2020.04.06.028001

**Authors:** Leo Chi U Seak, Konstantin Volkmann, Alexandre Pastor-Bernier, Fabian Grabenhorst, Wolfram Schultz

## Abstract

Rewarding choice options typically contain multiple components, but neural signals in single brain voxels are scalar and primarily vary up or down. In a previous study, we had designed reward bundles that contained the same two milkshakes with independently set amounts; we had used psychophysics and rigorous economic concepts to estimate two-dimensional choice indifference curves (IC) that represented revealed stochastic preferences for these bundles in a systematic, integrated manner. All bundles on the same ICs were equally revealed preferred (and thus had same utility, as inferred from choice indifference); bundles on higher ICs (higher utility) were preferred to bundles on lower ICs (lower utility). In the current study, we used the established behavior for testing with functional magnetic resonance imaging (fMRI). We now demonstrate neural responses in reward-related brain structures of human female and male participants, including striatum, midbrain and medial orbitofrontal cortex that followed the characteristic pattern of ICs: similar responses along ICs (same utility despite different bundle composition), but monotonic change across ICs (different utility). Thus, these brain structures integrated multiple reward components into a scalar signal, well beyond the known subjective value coding of single-component rewards.

**Significance Statement:** Rewards have several components, like the taste and size of an apple, but it is unclear how each component contributes to the overall value of the reward. While choice indifference curves of economic theory provide behavioural approaches to this question, it is unclear whether brain responses capture the preference and utility integrated from multiple components. We report activations in striatum, midbrain and orbitofrontal cortex that follow choice indifference curves representing behavioral preferences over and above variations of individual reward components. In addition, the concept-driven approach encourages future studies on natural, multi-component rewards that are prone to irrational choice of normal and brain-damaged individuals.

## Introduction

In daily life, we choose between options that have multiple components. In a restaurant, we can get, for the same price, a small but tasty steak or a larger but less tasty steak. In choosing the latter, we give up some taste for more meat. Or the components can be distinct objects, like a meal with small lasagne and big salad, or a meal with large lasagne and small salad; in choosing the latter, we give up some salad for more lasagne. In both cases, our preference for an option (steak or meal) is based on more than one component. To understand such choices, we need to know whether the value integrated from different components can be represented by scalar measures of preferences and their neuronal processes.

Functional magnetic resonance imaging (fMRI) studies investigated choices between bundles with multiple-components. Several brain regions are involved in such choices, including striatum (Hunt et al. 2014), frontal cortex (Hunt et al. 2014; Kurtz-David et al. 2019; Busemeyer et al. 2019), cingulate cortex (Kurtz-David et al. 2019; Busemeyer et al. 2019; Fujiwara et al., 2009) and insula (Busemeyer et al. 2019). One study showed encoding of values of gift cards that contained an amount component and a quality component (de Berker et al. 2019); other studies investigated irrational choices with monetary-gamble components (Kurtz-David et al. 2019) and addressed irrational attraction and decoy effects (Chau et al. 2014; Gluth et al. 2017; Chung et al. 2017). Whereas these studies demonstrated neural signals for multi-component rewards, they did not specifically investigate whether the signals captured the reward value integrated from multi-dimensional vectorial choice options. To resolve the issue would require to study how the increase of one component compensates for the decrease of the other component without changing the preference, and how such a trade-off is represented in scalar neural signals.

This trade-off mechanism constitutes the heart of indifference curves (IC) underlying Revealed Preference Theory (Samuelson 1938). Each two-component choice option is graphically represented at a specific x-y coordinate of a two-dimensional plot (Mas-Colell et al. 1995). All bundles that are equally preferred to each other (choice indifferent, indicating same utility despite different bundle composition) are located on the same IC irrespective of underlying variation in bundle composition. Preferred bundles are located on higher ICs (farther away from the origin, higher utility). This scheme is widely used for conceptualizing economic preferences in economics textbooks, consumer choice (Simonson 1989; Tversky & Simonson 1993; Rieskamp et al. 2006), animal choice (Kagel et al. 1975; Pastor-Bernier et al. 2017) and neuronal reward signals in animals (Pastor-Bernier et al. 2019). The preference scheme has been extended to stochastic choice (McFadden & Richter 1990; McFadden 2004), which is helpful for multi-trial statistical analyses of human brain responses. Thus, the question for the current study arises: would human blood-oxygen-level-dependent (BOLD) signals follow the characteristics of ICs that define the emergence of scalar measures from vectorial bundles?

We investigated scalar BOLD signals for two-component milkshakes with sugar and fat components that elicit subjective valuations and neural reward signals (Grabenhorst et al. 2010; Zangemeister et al., 2016). We used three revealed preference levels (three ICs, different utility), each estimated from five equally preferred bundles (indifference points, IPs, located on same IC, same utility despite different bundle composition). Participants were presented with choice options that contained one fatty and one sugary milkshake with specific amounts. We estimated psychophysical indifference points (IP) at which a Reference bundle and a Variable bundle were chosen with equal probability. From these IPs, we estimated well-ordered and non-overlapping ICs. Using two independent general linear models, we found that scalar BOLD responses in striatum, midbrain and medial orbitofrontal cortex followed the IC scheme: the responses varied monotonically across ICs but changed only significantly along individual ICs, indicating orderly integration of multi-component choice options into single-dimensional measures. The behavioral results of this study have been published in detail (Pastor-Bernier et al. 2020).

## Materials and Methods

### Participants

A total of 24 participants (19-36 years old with mean age 25.4 years; 11 males, 13 females) performed a binary choice task that was followed, in 50% of trials, by a Becker–DeGroot–Marschak (BDM) task inside the fMRI scanner using sugary and fatty milkshakes. All participants had known milkshake appetite, and none had diabetes or lactose intolerance. All participants provided written consent based on an information sheet. The Cambridgeshire Health Authority (Local Research Ethics Committee) approved this study. The behavioral results have been published with more details separately (Pastor-Bernier et al. 2020).

### Experimental design

The fundamental notion underlying this experiment posits that choice options consist of at least two components, and that preferences are revealed by observable choice. The multi-component choice options are called bundles. It is immaterial for the general concept of multi-component choice whether the individual components are parts of a single object (like size and taste of a steak in the example above) or constitute separate objects within a choice option (like lasagne and salad). Decision makers prefer bundles with larger or better components to those with smaller or worse components. Importantly, however, their preferences concern all components and are not directed at a single component alone. This property is manifested when participants prefer bundles in which one of the components of the preferred bundle is smaller than the same component in the non-preferred bundle (and the other component is large enough to overcompensate). At one point, participants may express equal preference for bundles in which the lower amount of one component is fully compensated by the higher amount in the other component, leading to choice indifference. We repeatedly measured choices with two options, each of which contained two milkshake components; the milkshakes constituted rewards, as shown by the voluntary consumption in all participants.

#### Stimuli and rewards

In each of the two bundles, we used stimuli to show the two milkshake components and their payout amounts (Fig. 1A). In each bundle stimulus, there were two rectangles aligned vertically. Each bundle component was indicated by the color of each rectangle. We extensively piloted various liquidized foods and liquids, and we found that milkshakes with a controlled mixture of fat and sugar give the most reliable across-participant behavioral performance. The presently used milkshakes with sugar and fat components that were found in previous studies to elicit subjective valuations and activate neural reward structures (Grabenhorst et al. 2010; Zangemeister et al., 2016). We delivered the milkshakes separately with a 0.5 s interval (see below). As drinks consisting of only sugar or only fat were considered as too unnatural, we used a high-fat low-sugar milkshake (75% double cream and 25% whole milk, with no sugar) as component A (top, blue), and a high-sugar low-fat milkshake (skimmed milk with 10% sugar) as component B (bottom, red). Inside each rectangle, the vertical position of a bar indicated the component’s physical amount (higher was more). We delivered the milkshakes to the participants using a custom-made silicone tubing syringe pump system (VWR International Ltd). The pump was approved for delivering foodstuffs and was controlled by a National Instruments device (NI-USB-6009) via the Data Acquisition Toolbox in Matlab. We displayed stimuli to participants and recorded behavioral choices using the Psychtoolbox in Matlab running on a Windows (Dell) computer (Pastor-Bernier et al. 2020).

**Figure 1.**
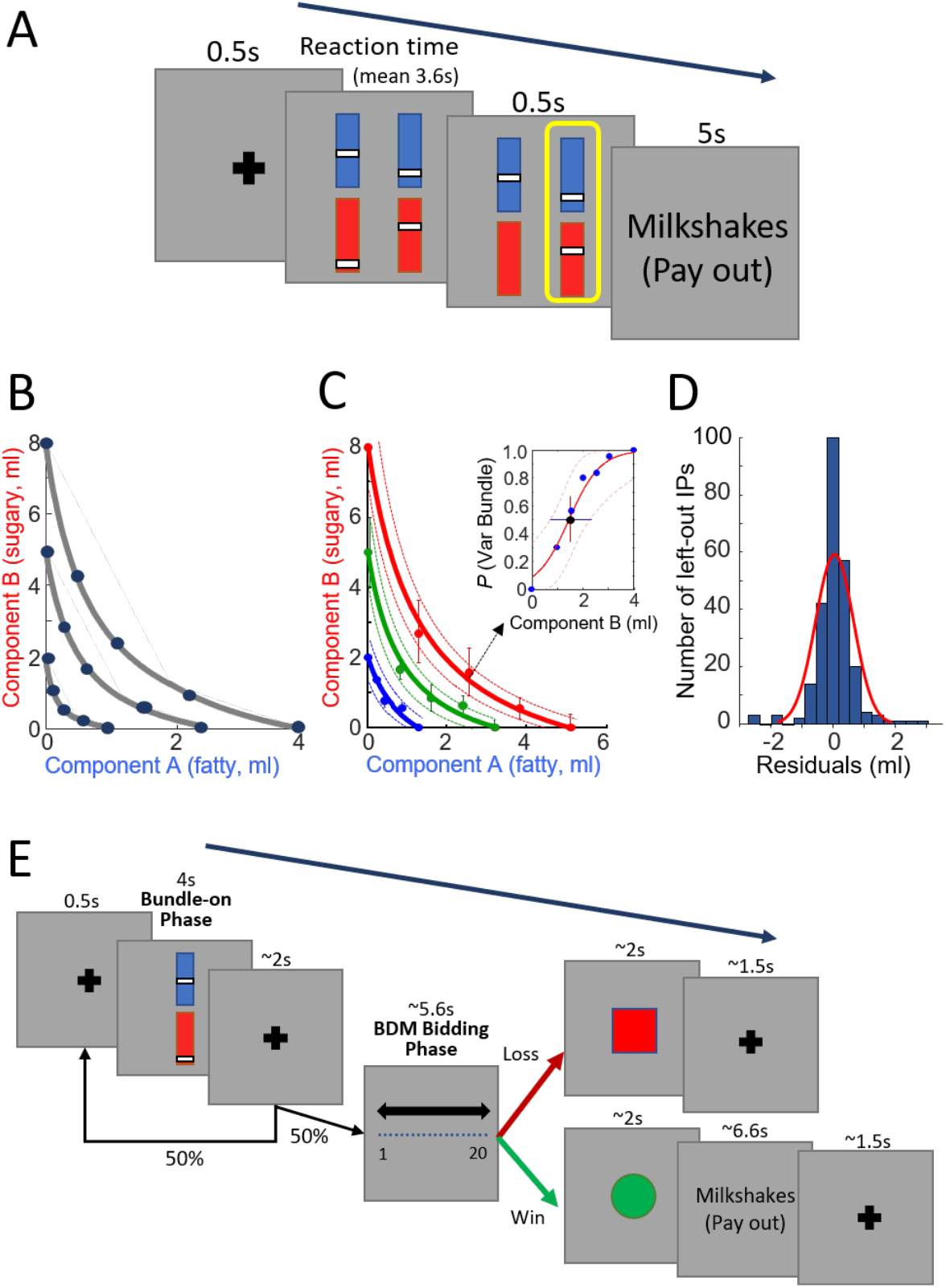
Experimental procedure and behavior. (A) Choice task outside the fMRI scanner. The participant chose between a reference bundle and a varied test bundle. Each bundle consisted of two components, Component A (blue bar) and Component B (red bar). The amount of each component was indicated to the participant by the height of a white bar (higher was more). Component A was a low-sugar, high-fat milkshake. Component B was a low-fat high-sugar milkshake. The two milkshakes of the chosen bundle were delivered at the end of each trial with a probability of P = 0.2. (B) Schematic diagram of three indifference curves (ICs) and five indifference points (IPs) on each IC (same data points as shown in Fig. 1F of Pastor-Bernier et al. 2020). (C) Example ICs from a typical participant. Solid lines represent three ICs (hyperbolically fitted by IPs). Dotted lines represent 95% confidence interval of the hyperbolic fit. The inset shows the psychophysical function of one IP. The IP (black dot in the inset) was estimated by probit regression on the test points (blue dot in the inset). The same graph is shown as Fig. 2A of Pastor-Bernier et al. (2020). (D) Histogram of residuals between fitted ICs (with a leave-one-out procedure) and left-out IPs across all participants. The residuals formed a normal symmetric distribution (red line). (E) Bundle task inside the fMRI scanner. At 4 s after the bundle-on phase, the participant performed in pseudorandomly selected 50% of trials an additional Becker-DeGroot-Marschak (BDM) task against the computer (bidding 1 - 20 UK pence). The reward was given if the participant won the BDM (bid ≥ computer bid).

#### Binary choice task before fMRI scanning

In the binary choice task, each participant revealed one’s preference in repeated choices between two bundle stimuli, each indicating the amounts of two milkshake components (Fig. 1A). The two bundles (stimuli) appeared on a computer screen simultaneously in front of the participant. The left and right positions of the bundles were fixed but pseudorandomly alternated. Each bundle stimulus included the same two kinds of milkshakes with independent physical amounts. Both stimuli appeared after a pseudorandomly varying interval (mean 0.5 s) after a central fixation cross. In each trial, the participant chose between the two bundles by pressing a button once (on a computer keyboard; left or right arrow corresponding to choosing left or right bundle). We defined reaction time as the interval between appearance of the two bundle stimuli and the participant’s button press. We delivered the two milkshakes to the participant from the chosen bundle with a probability *P* = 0.2 using a Poisson distribution; i. e. the milkshake combination of one out of an average of five chosen bundles was delivered, and no milkshake was delivered in the remaining trials. Component B (high-sugar low-fat milkshake) was delivered at a constant interval of 0.5 s after component A (high-fat low-sugar milkshake). We used this constant delay, instead of simultaneous delivery of two milkshakes or a pseudo-randomly alternating milkshake sequence, to prevent uncontrolled milkshake interactions, to maintain distinguishability of the individual milkshake rewards and to keep temporal discounting constant. Therefore, the utility of component B derived from both milkshake rewards and the temporal discounting specific for each milkshake. While the interval of 0.5 s was sufficiently short to not disrupt task performance and data collection, it was too short to completely prevent the high-fat milkshake blending into the subsequent high-sugar milkshake inside the participant’s mouth. As the interval was kept constant in all participants and at all times, the mixture provided a constant gustatory experience. Participants were asked not to eat or drink anything at least four hours before the task performance. However, satiety may still be a concern given the high fat and sugar content of our milkshakes. To address this issue, we set the probability of *P* = 0.2 payout schedule, limited each payout to 10.0 ml at most, and delivered no more than a total of 200 ml of liquid to the participant in a session. We addressed the issue with additional analyses and failed to find differential, sensory-specific satiety noticeable in choice probability measures (see below; Pastor-Bernier et al. 2020).

#### Psychophysical assessment of indifference points (IPs)

We used a psychophysical staircase method (Pastor-Bernier et al. 2020; Green, & Swets, 1966) to estimate the indifference points (IPs) at which, by definition, each of the two bundle options was chosen equally frequently (i.e. probability *P* = 0.5 for each option), indicating choice indifference for the options. We established bundles at 15 IPs for each participant and used them in the subsequent fMRI experiment.

To start the psychophysical procedure, we first set component A to 0 ml and component B to either 2 ml, 5 ml or 8 ml in the Reference Bundle. We then systematically varied the Variable Bundle. In the Variable Bundle, we first set the amount of its component A to one unit higher (mostly 0.5 ml, 1.0 ml or 2.0 ml); we thereby specified the amount of component A gained by each participant from the choice. We then randomly selected (without replacement) one amount of component B from a total of seven fixed amounts (multiples of 0.5 ml), which span the whole, constant range of amounts being tested. We repeatedly selected the amounts until we tested each of the seven amounts once. We repeated estimation for each IP six times using a sigmoid function (see Eqs. 1, 1a below), requiring a total of 42 choices for estimating each IP. The amount of component B in the Variable Bundle was usually lower than the one in the Reference Bundle at the IP. With these procedures, we assessed how much of component B a participant was willing to trade-in for an additional unit of component A.

We obtained more IPs from the participants’ choices between the fixed Reference Bundle and the Variable Bundle, in which the amount of component A was increased stepwise, at each step varying the amount of component B to estimate the choice indifference point at which the animal was indifferent between the two bundles. Thus, bundle position advanced from top left to bottom right on the two-dimensional IC (Fig. 1B). We are aware that testing with unidirectional progression may cause particular variations in IP estimations than testing in a random sequence or in opposite directions (Knetsch, 1989). However, our primary interest in this study was to investigate basic neural processes in close relation to unequivocally estimated IPs and ICs rather than addressing the more advanced features of irreversibility or hysteris in ICs.

We used three different fixed amounts of component B for the Reference Bundle (2 ml, 5 ml, or 8 ml), to obtain three IC levels. We estimated four IPs, together with the fixed reference bundle as an IP, at each of three indifference curves (ICs; i.e. revealed preference levels), resulting in 15 IPs, in a total of 504 choices (trials) among 84 different choice option sets in each participant (6 repetitions for 7 psychophysical amounts at each of the 12 IPs).

### Statistical analysis

#### Numeric estimation of indifference points

We used a sigmoid fit to numerically estimate the choice IPs. The fit was obtained from the systematically tested choices with a generalized linear regression. The generalized linear regressions used the *glmfit* function in Matlab (Matlab version R2015b) with a binomial distributed probit model, which is an inversed cumulative distribution function (G). More specifically, we apply the link function to the generalized linear regression y = β_0_ + β_1_B_var_ + ε and write it as:

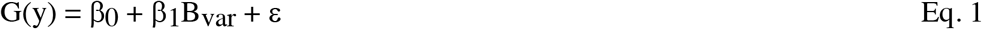

where y represents the number of trials the Variable Bundle is chosen in each block of a six-repetition series, β_0_ represent the constant offset, β_1_ represent the regression slope coefficient, B_var_ represent the physical reward amount (ml) of component B in the Variable Bundle, and ε represent the residual error. We used the probit model as it assumes a multivariate normal distribution of the random errors, which makes the model attractive because the normal distribution gives a good approximation to most of the variables. The model does not hypothesize error independence and is frequently used in econometrics (Razzaghi, 2013). On the other hand, the logit model, which is also commonly used in economics, is simpler to compute but has more restrictive hypotheses on error independence. Our preliminary data had shown a similar fit for both the logit and probit model, therefore, we used the probit model fit because of its less restrictive hypotheses. Thus, we approximated the IPs with the probit-model sigmoid fit, which can be written as follows:

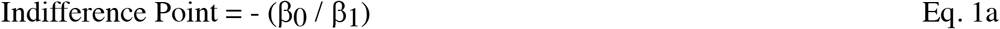

where β_0_ and β_1_ represent coefficients of the generalized linear regression (Eq. 1). We obtained these coefficients from the probit analysis (Amemiya, 1981).

#### Indifference curves (ICs)

In each participant, we obtained each single IC separately from an individual set of five equally revealed preferred IPs with differently composed bundles using a weighted least-square non-linear regression. We used a weighted regression to account for choice variability within participant; the weight was defined as the inverse of the standard deviation of the titrated physical amount of component B at the corresponding IP (the IP having been estimated with the probit regression). We estimated the best β coefficients from the least-square regression to obtain a single IC (utility level), using the basic hyperbolic equation:

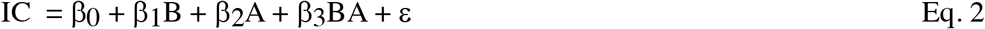

where A and B represent physical amounts of component A and component B (ml), which refer to the x and y axis, respectively. Note that (β2 / β1) is the slope coefficient and β_3_ is the curvature coefficient of the non-linear least-square regression. As IC is a constant (representing one utility level), we merged the IC constant with the offset constant (β_0_) and the error constant (ε) into a common constant k. To draw the ICs, we calculated the amount of component B from the derived equation as a function of the amount of component A:

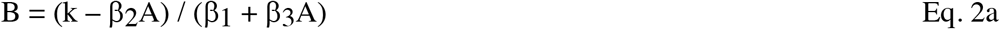

We graphically displayed the fitted ICs (Fig. 1B, C) by plotting the pre-set physical amount of component A as the x coordinates, and calculated the fitted amount of component B, based on Eq. 2a, as the y coordinates. We estimated the error of the hyperbolic fit as the 95% confidence interval. When calculating the ICs, we gave less weight to the IP with higher error. This model offered good fits in our earlier work (Pastor-Bernier, et al. 2017; 2019; 2020). In this way, five IPs aligned to a single fitted IC. For each participant, we fitted three ICs representing increasing revealed preference levels (low, medium, high) farther away from the origin (Fig. 1B, C). The indifference map that resulted from the 3 x 5 IPs was unique for each of the 24 participants. The indifference maps of the 24 participants were presented before (Pastor-Bernier et al. 2020).

#### Leave-one-out validation of ICs

We used a leave-one-out analysis to test the validity of the hyperbolic IC fit to the IPs. We systemically removed one IP in each IC (excluding the initial Reference Bundle at x = 0), and then fitted the IC again using the hyperbolic model. We then assessed the differences (deviation) between the original IC (without IP removal) and the new IC without the one left-out IP. The deviation was defined as the Euclidean distance of component B between the original (left-out) IP and the IP estimated from the refitted IC:

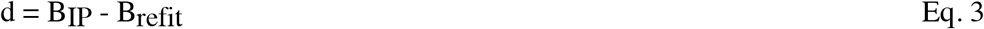

with d representing the difference (i.e. residual; in ml; y-axis), BIP representing the physical amount of component B in the left-out IP (ml), and B_refit_ representing the estimated physical amount of component B in the refitted IC (ml). In this way a residual of 0 ml suggested that removal of the left-out IP did not change the shape of that IC, while any residual unequal to 0 ml could quantify the deviation.

#### Control of alternative choice factors

To assess the potential influence of other factors affecting the participants’ choice, we performed a logistic regression fit on choices to test whether the choices were indeed explained by the bundle components. We performed a random-effect logistic regression on the choice data from each participant as follows:

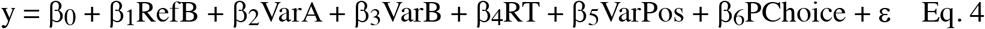

with y as a dummy variable (either 1 or 0, indicating choosing or not choosing the Variable Bundle), RefB as physical amount (ml) of component B in the Reference Bundle, VarA and VarB as physical amount (ml) of components A and B in the Variable Bundle, RT as reaction time (ms), VarPos indicating left or right position (0 or 1) of the Variable Bundle stimuli shown on the computer screen relative to the Reference Bundle, and PChoice representing choice of the previous trial (0 or 1). Each β coefficient was normalized by multiplying the standard deviation of the respective independent variable and dividing by the standard deviation of the dependent variable (y). We subsequently used a one-sample t-test against 0 to assess the statistical significance of each of the beta (β) coefficients.

We assessed the normalized beta (β) coefficients and p-values for each individual participant and then calculated averages across 24 participants. With the regression model, we found a negative correlation of choosing the Variable Bundle and the amount of component B in the Reference Bundle (RefB: *β* = −0.43 ± 0.16, *P* = 0.020 ± 0.005; mean ± SEM) (amount of component A in the Reference Bundle was always a constant 0 ml). We also found positive correlation of choosing the Variable Bundle and amount of both component A and component B in the Variable Bundle (VarA: *β* = 0.67 ± 0.16, *P* = 0.009 ± 0.004; VarB: *β* = 0.94 *β* 0.33, *P* = 0.012 ± 0.009). We further found that for these three variables, the beta (β) coefficients significantly differed from 0 with one-sample t-tests (*P* = 0.012, *P* = 0.00088 and *P* = 0.00028, respectively), confirming the robustness of these β. Thus, we confirmed that the choices depended on the amount of reward of both Variable and Reference Bundle. We also validated that both bundle components were important for the choices. All remaining variables in the regression, including reaction time, left or right position of the Reference Bundle on the computer screen and choice of the previous trial, failed to account significantly for the participant’s current choice (*P* = 0.754 – 0.988 ± 0.003 – 0.290). We therefore conclude that, in our experiment, the bundles with their two components, instead of other factors, account for the revealed preference relationships.

#### Satiety control

Besides considering other components in the design, we also tested potential effects of satiety. Satiety may have affected the preferences for the two bundle components, even if the rewards were paid out only in one fifth of the trials on average and were limited to less than 200 ml. Differences in devaluation between the two component milkshake might be a major factor for changing in an uncontrolled manner the currency relationship of the two components. This kind of unequal devaluation should result in a graded change in the instantaneous choice probability around the IPs over the test steps of 42 trials. We used the following equation to calculate the instantaneous choice probability:

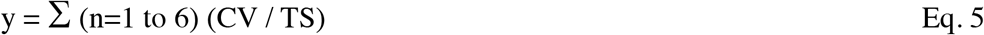

with y representing the instantaneous probability (*P* ranging from 0.0 to 1.0), CV represent choice or not-choice of Variable Bundle (1 or 0), and TS represent test step (repetition 1-6).

We found only insignificant fluctuations in choice probabilities, without any consistent upward or downward trend in the 1-way repeated measures ANOVA, together with the post-hoc Tukey Test (above IP: F (5, 41) = 0.28, *P* > 0.05; below IP: F (5, 41) = 1.53, *P* > 0.05).

### Behavioral task during fMRI scanning

During scanning, we used a value elicitation task that allowed more trials in a shorter time frame. At the beginning of each trial, one bundle was shown to the participant for 5 s (bundle-on phase in Fig. 1E) in the center of the computer monitor after the initial fixation period (500 ms). The bundle was pseudorandomly selected from the 15 IP bundles in three ICs of each participant. Bundle composition (amounts of the two components) was set in each participant according to performance in the binary choice task before fMRI scanning. Hence, the 15 bundles for each participant were not identical across participants. Subsequently, a fixation cross appeared for a pseudorandomly varying interval (mean 2s). In 50% of the trials (pseudorandomly selected), the task was terminated after this fixation cross.

In the other 50% of the trials, we presented the participant with a Becker-DeGroot-Marschak (BDM) task that was akin to a second price auction (Becker, DeGroot, & Marschak, 1964). This task served as an independent mechanism that related the estimated ICs to stated utility. In the BDM (bidding phase in Fig. 1E), we gave the participant a fresh 20 UK pence endowment on each trial. Using this endowment, the participant bid for a two-component bundle against a pseudorandom computer bid (extracted from a normal distribution with replacement). To bid, the participant moved a cursor, shown on the computer screen, horizontally with the left and right keyboard arrows. We registered the BDM bid (position of the cursor) 5 s after presenting the bidding scale to the participant. When bidding no less than the computer, the participant received the bundle (milkshake) reward from both components and paid the monetary value equal to the computer bid. By contrast, when bidding less than the computer, the participant lost the auction, paid nothing and would not get any bundle (milkshake) reward. We showed the participant the result of the auction immediately after having placed the bid, by displaying a respective win (green circle) or loss (red square) stimulus on the computer monitor (Fig. 1E); when winning the bid, the participant received the milkshake rewards in the sequence and frequency as in the binary choice task.

We first selected one bundle randomly (without replacement) from the participant-specific set of 15 bundles (the 15 bundle IPs used to fit the 3 ICs as shown in Fig. 1). Then we showed the participant the selected single bundle during the bundle-on phase. We presented each of the 15 bundles to the participant for 24 times, resulting in a total of 360 trials, which included 180 trials (50%) with BDM bidding (Fig. 1E), and we used the average of these bids as the participant’s BDM-estimated utility.

First, we assessed whether the BDM bids increased for bundles across revealed preference levels but were similar for IP bundles on the same revealed preference level, using Spearman rank correlation analysis and further confirmation with the Wilcoxon signed-rank test (note that this analysis used the coordinates of the individual IPs to which the ICs had been fitted, not the IC coordinates themselves). We also performed a generalized linear regression with a Gaussian link function (random-effect analysis) for each participant and then averaged the β coefficients and p-values across all participants. We used the following generalized linear regression:

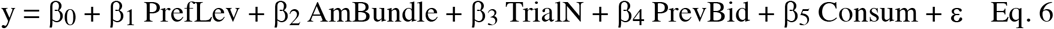

with y representing amount of monetary bid, PrefLev representing revealed preference level (low, medium, high), AmBundle representing the summed amount (ml) of component A and component B in the currency of component A (converted with Eq. 2a), TrialN representing trial number, PrevBid representing amount of monetary bid in the previous trial, and Consum representing accumulated consumption amount (ml) of component A and component B until that point in the experiment. We normalized each β coefficient by multiplying the standard deviation of the respective independent variable, and then dividing by the standard deviation of the dependent variable y. We performed a subsequent one-sample t-test against 0 to assess the significance of each beta (β) coefficient across all 24 participants. We found significant beta (β) coefficients of BDM monetary bids to the preference level (PrefLev: β-coefficient difference from 0: P = 0.000026 with one-sample t-test; mean across all 24 participants: β = 0.47 ± 0.09, P = 0.016 ± 0.015; mean± SEM) and bundle amount (AmBundle: P = 0.0278; β = 0.15 ± 0.13; P = 0.020 ± 0.017), but not in trial number (TrialN: β= −0.10 ± 0.25; P = 0.726 ± 0.354), previous trial bid (PrevBid: β= 0.12 ± 0.11; P = 0.676 ± 0.427) nor consumption history (Consum: β= 0.12 ± 0.11; P = 0.224 ± 0.185).

### fMRI data acquisition

The functional neuroimaging data in this study were collected using a 3T Siemens Magnetom Skyra Scanner at the Wolfson Brain Imaging Centre, Cambridge, UK. Echo-planar images (T2-weighted) with blood-oxygen-level-dependent (BOLD) contrast were acquired at 3 Tesla across two days with each participant. All images were in plane resolution 3 x 3 x 2 mm, 56 slices were acquired with 2 mm slice thinness, repetition time (TR) = 3 s, echo time (TE) = 30 ms, −90 deg flip angle and −192 mm field of view. To reduce signal dropout in medial-temporal and inferior-frontal regions during the scanning, the acquisition plane was tilted by −30 degrees and the z-shim gradient pre-pulse was implemented. We also applied MPRAGE sequences and co-registered to acquired high-resolution T1 structural scans for group-level anatomical localization with 1 x 1 x 1 mm^3^ voxel resolution, slice thickness of 1 mm, 2.3 s TR, 2.98 ms TE, 9 deg flip angle and 900 ms inversion time.

### fMRI data analysis

We used the Statistical Parametric Mapping package to analyze the neuroimaging data (SPM 12; Wellcome Trust Centre for Neuroimaging, London). We pre-processed the data by realigning the functional data to include motion correction, normalizing to the standard Montreal Neurological Institute (MNI) coordinate, and then smoothing using a Gaussian kernel with the full width at half maximum (FWHM) of 6 mm within data collected on the same day. We then segmented the data to extract white matter, grey matter and cerebrospinal fluid (CSF) and followed by co-registering the two-day data using the T1-weighted structural scans from each day. We then applied a high-pass temporal filter to it with a 128 s cut-off period. We applied General linear models (GLMs), which assumed first-order autoregressions, to the time course of activation. We modeled event onsets, in the time course of activation, as single impulse response functions convolved with the canonical hemodynamic response. We included the time derivatives in the functions set and defined linear contrasts of parameter estimates to test the specific effect in each participant’s dataset. We obtained voxel values for each contrast in the format of a statistical parametric map with corresponding t-statistic. We applied a standard explicit mask (mask_ICV.nii) at the first level analysis to mask out all activations outside of the brain. To test our specific hypotheses, we used the following GLMs:

#### General linear model 1 (GLM1)

This GLM served to search for regions whose stimulus-induced brain activations varied across ICs (high > low) but not along the same ICs in the bundle-stimulus-on phase (two-level t-test analysis, Fig. 2). For each participant, we estimated a GLM with the following regressors (R) of interest: (R1-R15) as indicator functions for each condition during the bundle-on phase (for the 15 different bundles), at the time when participant was presented with the visual bundle cue representing the milkshakes bundles; (R16) as indicator function for the BDM bid, at the time when the participant made the bid; (R17) as R16 that was modulated by the response to the participant’s bid (1 - 20); (R18) as indicator function for the losing bid, at the time when the participant was presented with visual cues showing the loss of bidding of the trial; (R19) as indicator function for the auction win phase, at the time when the participant was presented with the visual cues representing the winning of bidding; (R20) as indicator function for the reward phase, i.e. the times when participants received the milkshakes; (R21) as R20 that was modulated by reward magnitude (in mL). Regressors R16 - R19 were not used further for this analysis and served only to regress out potential BDM effects in the 50% of trials that included BDM.

**Figure 2.**
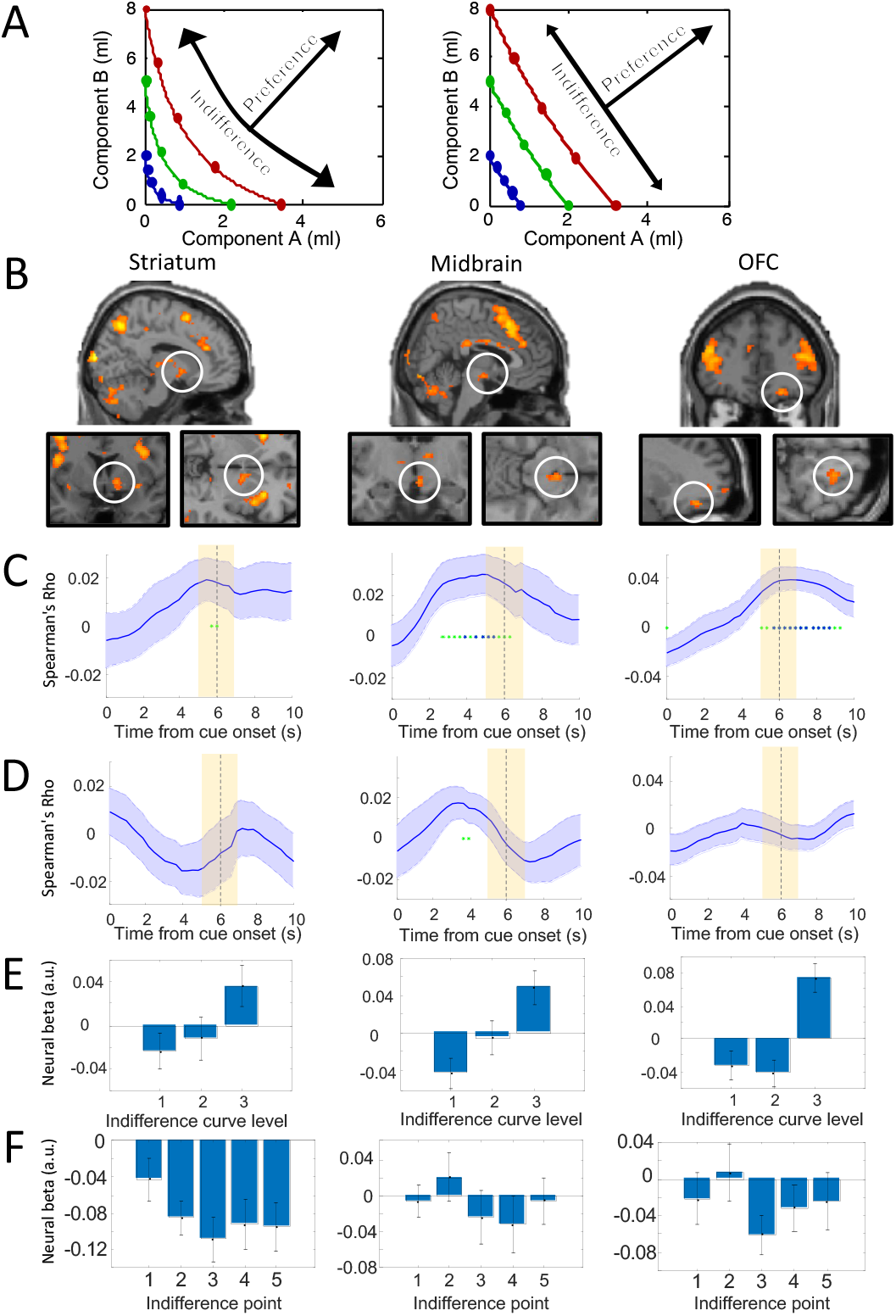
BOLD responses following the revealed preference scheme of two-dimensional indifference curves (ICs). (A) Schematic of the analysis method used in GLM1 (arrows): significant BOLD signal across ICs (increasing utility) but not within ICs (same utility despite different bundle composition, as inferred from choice indifference). Participants typically showed convex ICs (left) or linear ICs (right). (B) BOLD responses discriminating bundles between ICs (map threshold p < 0.005, extent threshold ≥ 10 voxels), but no discrimination between bundles along same ICs (map threshold p > 0.005; i.e. exclusive mask for brain response falls along the same ICs with threshold p=0.005) in a group analysis. For activations identified with F contrast, see Fig. 2-1. For activations identified with the lower threhold of p < 0.001, see Fig. 2-2. (C) Across-IC Spearman rank analyses of brain activations. The Rho coefficients followed the haemodynamic response function (HRF) across the 3 IC levels in the ROIs of the three brain structures shown above in B. Solid blue lines represent mean Rho from 24 participants; + SEM. Yellow shaded boxes show analysis time window. Green asterisks p < 0.05, blue asterisks p < 0.01 for t-test of Spearman’s Rho against zero. The BOLD responses (input of the Spearman rank analyses; with motion parameters regressed out) were extracted from the peak voxels of each participant using with a leave-one-out procedure (see Methods). (D) Along-IC ROI activations. The Spearman rank analyses indicated hardly any significance along same ICs in ROIs of the three brain structures shown in B. (E) Bar charts of neural beta coefficients of GLM1 for the three IC levels in the three brain structures shown in B in 24 participants. Bars show mean ± SEM. (F) Bar charts of neural beta coefficients of GLM1 for all five indifference points (IPs) on same IC levels (neural beta coefficients were averaged across the three IC levels in each participant) in 24 participants. Insignificant differences in one-way Anova: striatum: p = 0.3845, F(4,115) = 1.05, midbrain: p = 0.6828, F(4,115) = 0.57; OFC: p = 0.5672, F(4,115) = 0.74.

**Figure 2-1.**
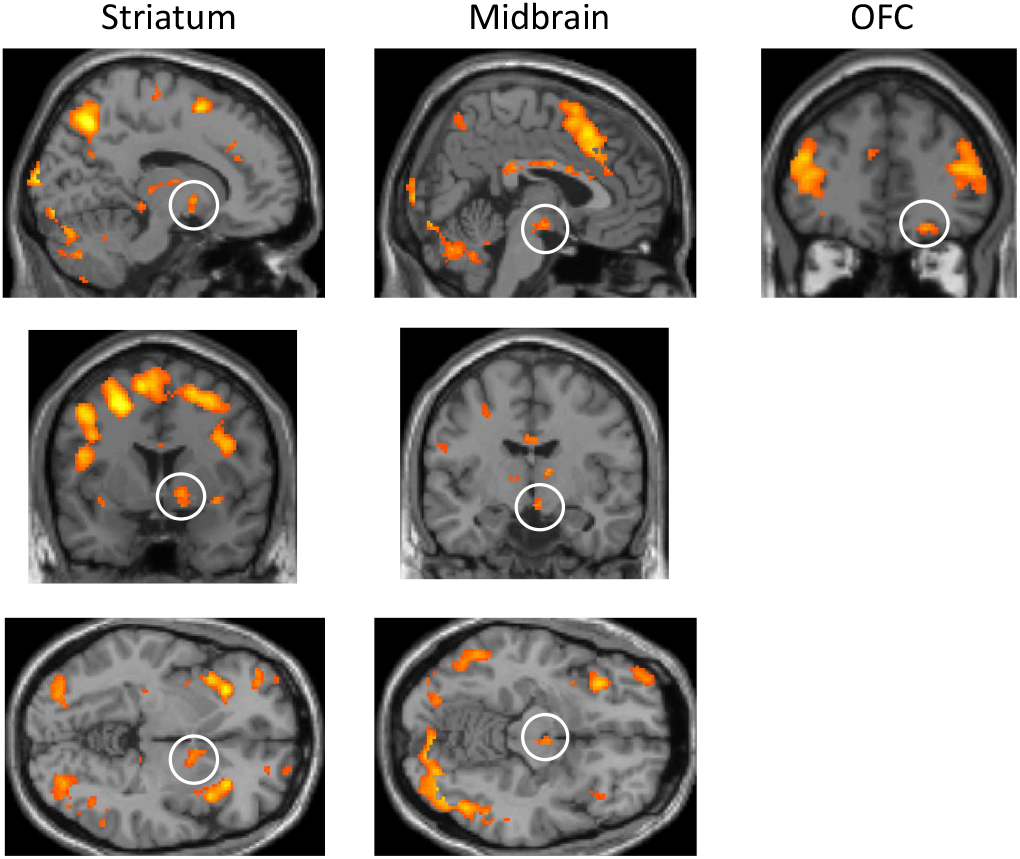
BOLD responses discriminating bundles between indifference curves (ICs) identified with F contrast (map threshold p < 0.005, extent threshold ≥ 10 voxels, high>low), but no discrimination between bundles along same ICs (map threshold p > 0.005; i.e. exclusive mask for brain response to bundles on same ICs with threshold p=0.005) in a group analysis. OFC: orbitofrontal cortex.

**Figure 2-2.**
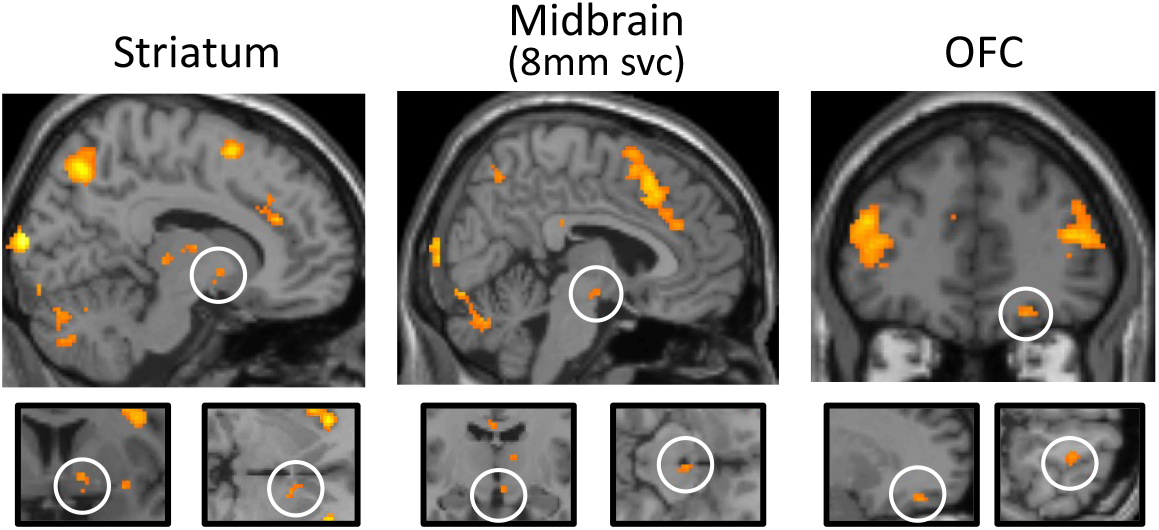
BOLD responses discriminating bundles between ICs with lower threshold (map threshold p < 0.001, extent threshold ≥ 10 voxels, high > low), but no discrimination between bundles along same ICs with T contrast (map threshold p > 0.005; i.e. exclusive mask for brain response to bundles on same ICs with threshold p=0.005) in a group analysis. Svc: small volume corrected.

In the second (group random-effects) level analysis, we entered the all 24 participant-specific linear contrasts of the first-level regressors R1-R15 (representing 5 bundles on each of the three preference levels) into t-tests (high > low revealed preference level) using Flexible Factorial Design, resulting in group-level statistical parametric maps. In the Flexible Factorial design matrix (second-level analysis), the following second-level regressors were used: (R1-24) indicator functions of participant’s identifier representing participant 1 - 24 (within participant effect); (R25-27) indicator functions of the three revealed preference levels (across ICs) (R28-32), indicator functions of the 5 bundles representing amount of Component A in increasing magnitude or amount of Component B in decreasing magnitude (along the same ICs). We first calculated the main contrast image based on high>low revealed preference level (t-tests). Second, we calculated a mask contrast based on 5 bundles of Component A in increasing magnitude (t-tests). Third, we calculated another mask contrast based on 5 bundles of Component B in increasing magnitude (t-tests). The final result of GLM1 was represented by the main contrast (high>low revealed preference level) masking out (with exclusive mask) the two mask contrasts, controlling of the brain responses along the same ICs.

#### General linear model 2 (GLM2)

This GLM identified regions associated with the binary comparisons of partial physical non-dominance bundles (Fig. 3). The GLM searched for brain regions in which activations were higher for bundles that were on a higher revealed preference level than bundles in which one component was physically higher than in the preferred bundle (partial physical non-dominance). In the first-level estimation, regressors were the same as in GLM1 with the 21 regressors described above. In the second-level analysis, we entered all pairs of bundles that met the following criteria: (Bundle 1): partial physical non-dominance bundles with higher revealed preference level, but less (with at least 0.2mL less in Components A or 0.4mL less in Component B) in one component; (Bundle 2): partial physical dominance bundles with lower revealed preference level, but more in one component. A third level group-level analysis (one-sample t-test) was performed with contrast images from the second level to generate group-level statistical parametric maps across 24 participants.

**Figure 3.**
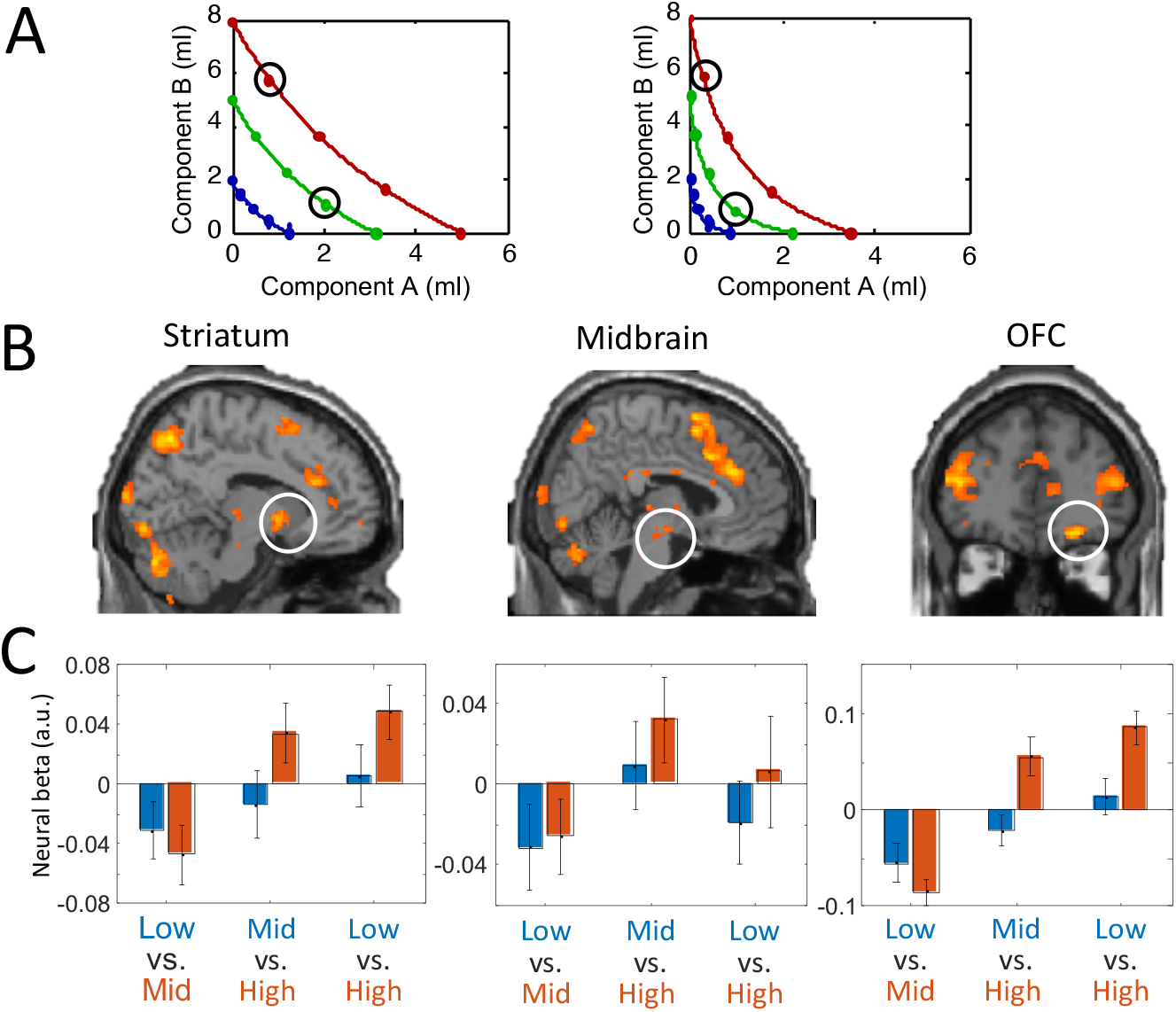
Higher BOLD responses to more preferred (but physically partially dominated) bundles positioned on different indifference curves (ICs). (A) Two examples of binary bundle comparison. Each pair of black circles indicates one binary comparison in one participant. (B) Brain regions activated more by preferred bundles compared to alternative bundles in group analysis with GLM2. Map threshold p < 0.005, extent threshold ≥ 10 voxels. For activations identified with the lower threshold of p < 0.001, see Fig. 3-1. (C) Bar charts showing neural beta coefficients of regression in ROIs of three brain structures in the population of 24 participants. Each group of bars (3 groups in each ROI) shows the beta coefficients for bundles in partial physically dominating relationships on different ICs: low vs. mid; mid vs. high and low vs. high. Orange bars represent the higher preference level and blue bars represent the lower preference level in each comparison. The bars show the mean ± SEM. For activations at peak voxels, see Fig. 3-2.

**Figure 3-1.**
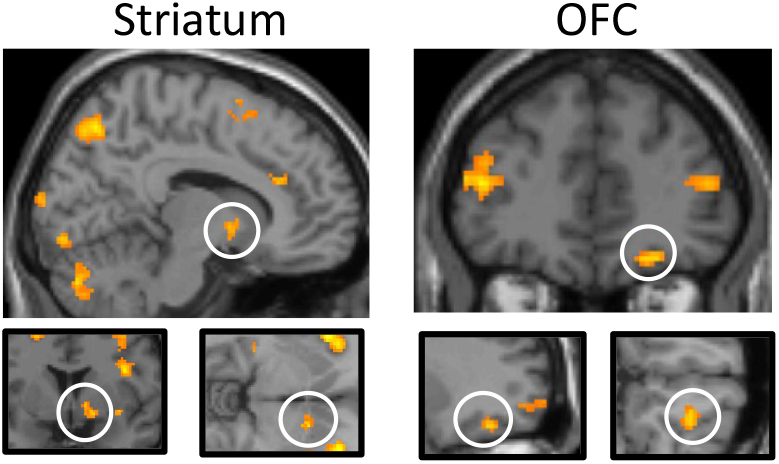
Higher BOLD responses to more preferred (but physically partially dominated) bundles positioned on different indifference curves with stricter thresholds (Map threshold p < 0.001, extent threshold ≥ 10 voxels) in striatum (left) and OFC (right).

**Figure 3-2.**
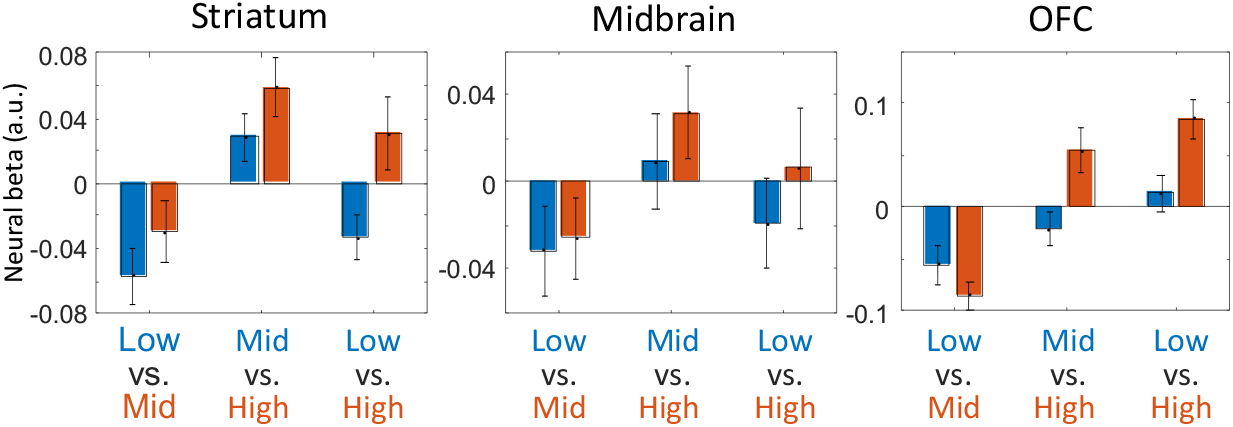
Bar charts showing neural beta coefficients of regression at peak voxels in ROIs (with ROIs coordinate extracted from GLM1 using leave-one-out procedure) of three brain structures in the population of 24 participants. Each group of bars (3 groups in each ROI) shows the beta coefficients for bundles in partial physically dominating relationships on different indifference curves (IC): low vs. mid; mid vs. high and low vs. high. Orange bars represent the higher preference level and blue bars represent the lower preference level. The bars show the mean ± SEM.

#### General linear model 3 (GLM3)

This GLM identified brain regions in which activity correlated with the amount of BDM bid (0 - 20 pence) during the bidding phase (Fig. 4B). In the first-level estimation, we used the following regressors and parametric modulators: (R1) as indicator function of bundle-on phase; (R2) as R1 modulated by amount of BDM bid; (R3) as indicator function of BDM bidding phase (50% of trials); (R4) as R3 modulated by amount of BDM bid; (R5) as indicator function of intertrial interval when there was no bidding phase (50% of trials); (R6) as indicator function at onset of the loss cue, when the participant lost the BDM bidding; (R7) as indicator function at onset of the win cue, when the participant won the BDM bidding; (R8) as indicator function at onset of milkshake delivery; (R9) as R8 modulated by physical amount of milkshake; (R10) as contrast of win cue onset versus loss cue onset; (R11) as contrast of loss cue onset versus win cue onset. In the second-level analysis, a one-sample t-test analysis was performed with contrast images from the first level to generate group-level statistical parametric maps across 24 participants.

**Figure 4.**
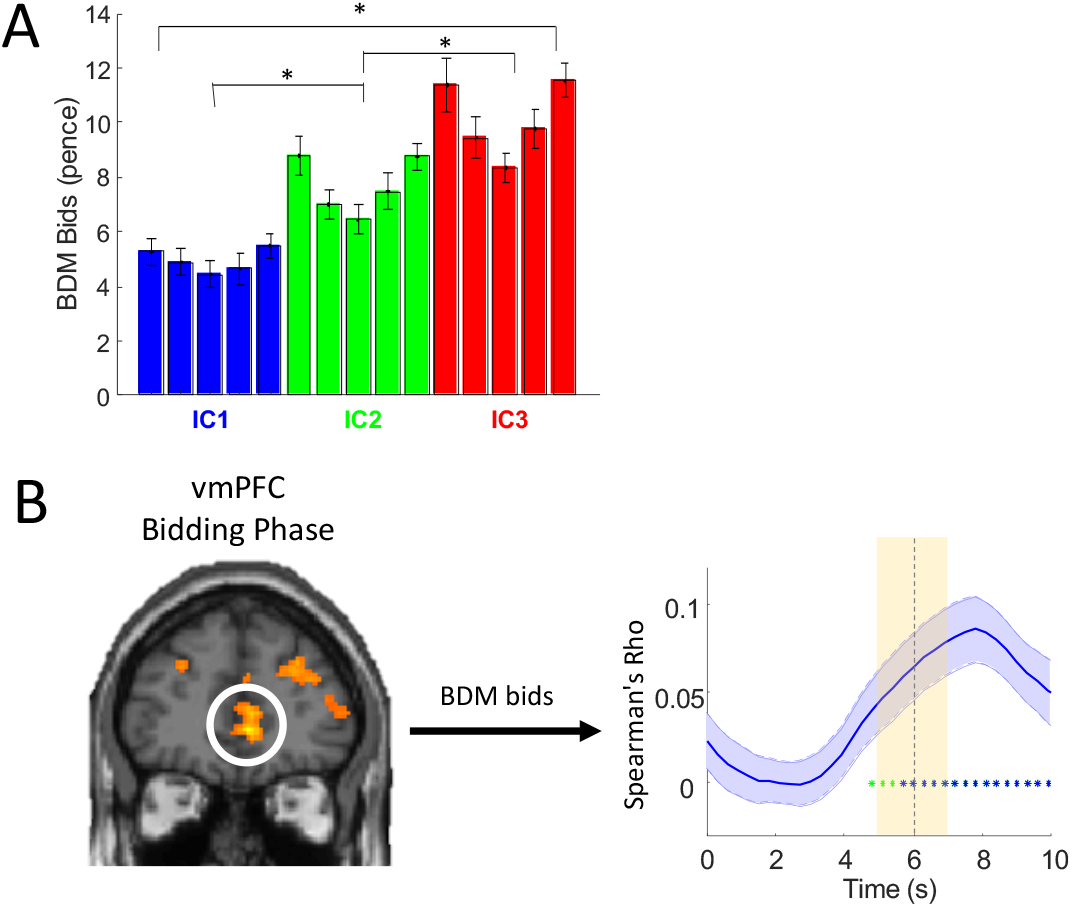
Activation in ventromedial prefrontal cortex (vmPFC) during BDM bidding. (A) Bar chart for BDM bids for 15 bundles of 24 participants (mean ± SEM). The colors of the bars indicate the indifference curves (IC) to which the bundle belongs (blue = low IC; green = middle IC; red = high IC). Spearman rank correlation: across ICs: Rho = 0.5710, p = 1.5659 x 10^−32^; within IC: Rho = 0.0219, p = 0.6791. Wilcoxon signed rank test: IC1 vs. IC2: p = 1.2802 x 10^−20^; IC2 vs. IC3: p = 8.0748 x 10^−21^; IC1 vs. IC3: p = 1.5954 x 10^−19^. (B) vmPFC activation during bidding phase (GLM3: activation correlated with BDM bids; threshold p < 0.005, extent threshold ≥ 10 voxels). Spearman rank analysis (right) showed significant Rho coefficient across bids. For additional activation in dorsal striatum, see Fig. 4-1.

**Figure 4-1.**
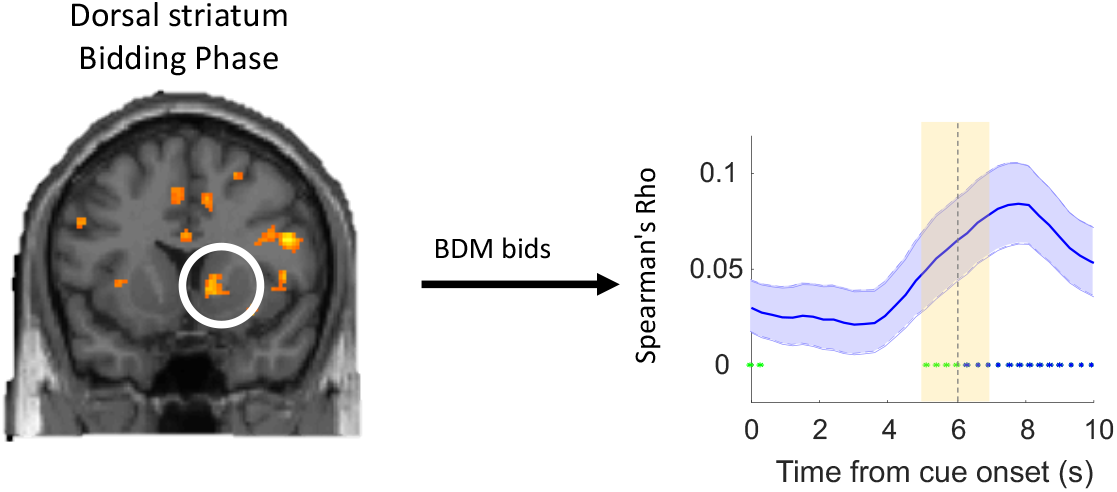
Dorsal striatum activation during bidding phase (GLM3: activation correlated with the amount of BDM bids; threshold p < 0.005, extent threshold ≥ 10 voxels). Brain map (left) shows dorsal striatum activity during bidding phase. Spearman rank analysis (right) showed significant Rho coefficient across bids during bidding phase in dorsal striatum.

#### Small volume corrections

To derive coordinates for small-volume correction in GLM1 and GLM2, we entered the term “reward anticipation” in the Neurosynth meta-analysis database (Yarkoni et al., 2011) to obtain MNI coordinates. The meta-analysis employed a total of 92 independent studies that showed correlation of value elicitation with various brain regions. Our study used MNI coordinates of ventral striatum [12, 10, −8], medial orbital frontal cortex (mid-OFC) [20, 46, −18] and midbrain [8, −18, −14], obtained from this Neurosynth meta-analysis database. We used a sphere with 6 mm radius for midbrain and striatum, and 10 mm for OFC, following the common approach of using 6 mm radius spheres for subcortical structures and larger spheres for cortical structures (Zangemeister et al., 2016, De Martino et al., 2009, Chib et al., 2009).

We aimed at finding activity correlating with the BDM bid in GLM3. Therefore, for small volume correction analysis in GLM3, we used a MNI coordinate of dorsal striatum [12, 14, 4] found in a previous study with BDM bidding (De Martino et al., 2009). We did not use coordinates from Neurosynth in GLM3 because datasets related to BDM or other auctions were not available in the Neurosynth database.

### Region-of-interest (ROI) analysis

We selected significantly activated regions from brain maps established with GLM1, GLM2 or GLM3 for further ROI analysis. We extracted raw BOLD data from ROI coordinates based on group clusters, which we defined independently for each participant using a leave-one-out procedure based on the result of GLM1, GLM2 or GLM3. In the leave-one-out procedure, we re-estimated the second-level analysis 24 times, each time leaving out one participant, to define the ROI coordinates for the left-out participant. Following data extraction, we applied a high-pass filter with a cut off period of 128 s. The data was then z-normalized, oversampled by a factor of 10 using the Whittaker–Shannon interpolation formula, and separated into trials to produce a matrix of trials against time.

A total of 3 ROI analyses were performed in this study. First, a Spearman rank analysis was used to examine BOLD signals that changed across ICs but not along ICs (corresponding to GLM1 and GLM3). Second, a bar chart was used to illustrate the three revealed preference levels in different ROIs (corresponding to GLM1). Third, a bar chart was used to show activation changes between bundles with partial physical non-dominance on different revealed preference levels (corresponding to GLM2).

#### Spearman rank

In the Spearman rank analysis, we first regressed out the motion parameters (artefact) from the BOLD response with generalized linear models. Then we used the participant’s residual BOLD response to generate time courses of Spearman rank correlation (Rho) coefficients.

For GLM1, we tested the correlation between BOLD response (during the bundle-on phase) and revealed preference level (across-IC analysis). We then calculated group averages and standard errors of the mean for each time point for all participants, yielding averaged participant effect size time courses (Fig. 2C). In the along-IC analysis, we ranked the bundles along the same IC with individual participant’s BOLD signal (Fig. 2D). A subsequent one-sample t-test against 0 served to assess the significance of the Rho coefficients across subjects.

For GLM3, we tested the correlation of the BOLD response (BDM bidding phase) and the amount of BDM bids. Similar to GLM1, we then calculated group averages and standard errors of the mean of the Rho coefficients for each time point for all participants (Figs. 4B, 4-1). A subsequent one-sample t-test against 0 served to assess coefficient significance.

#### Bar chart for revealed preference level analysis

We used bars to illustrate how different IC levels were encoded in each region of the brain. To generate an ROI bar chart, the BOLD response was first extracted using the leave-one-out procedure described above. For each participant, we obtained three generalized linear model fits to the BOLD signal at timepoint 6 s. In each generalized linear model fit, the identifier of one level of revealed preference was entered as a regressor (dummy variable, e. g. 1 for bundles with high preference level and 0 for middle or low preference level) together with motion parameter regressors, which served to eliminate the motion artefact. We obtained beta (β) coefficients of each level of revealed preference from the fit and then calculated the mean and standard error of the beta coefficient. We then plotted the bar charts shown in Fig. 2E. Paired t-tests were used to compare beta coefficients between different revealed preference levels. As a control, we also obtained beta (β) coefficients of 5 indifference points from the same level of revealed preference, averaged across the three levels, and then calculated the mean and standard error of the beta coefficient across participants. We then plotted the bar charts shown in Fig. 2F. One-way ANOVAs were used to compare beta coefficients between the 5 indifference points.

#### Bar chart for partial physical non-dominance analysis

A bundle was defined as being partially physically non-dominant over another bundle if one of its milkshake components had a physically lower amount than the same component in the dominated bundle. Thus, the revealed preferred bundle was partially physically non-dominant. For an ROI analysis of partial physical non-dominance, we fitted three generalized linear models to the BOLD response with bundle identifiers, which were two dummy variables representing partially physically dominance bundles (lower revealed preference despite larger physical amount in one milkshake) and partial physical non-dominance bundles (higher revealed preference despite smaller physical amount in one milkshake). Three generalized linear models were used to fit bundles in low vs. middle, middle vs. high, and low vs. high comparisons, respectively. The domination was defined as at least 0.2 ml more for component A or at least 0.4 ml more for component B, as in GLM2. We calculated the mean and standard error of the averaged beta (β) coefficients across participants at time point 6 s and plotted the bar chart as shown in Fig. 3C. Paired t-tests were used to compare beta coefficients between partial physical dominance bundles and partial physical non-dominance bundles. Motion parameters were also used as regressors for each participant to eliminate motion artefacts. In addition to extracting BOLD signal with leave-one-out peaks of GLM2 (Fig. 3C), we also extracted BOLD signal with leave-one-out peak from GLM1 (Fig. 3-1) to confirm the robustness of this analysis.

### Reward prediction errors (RPE)

The current task did not involve learning in which reward would occur in a partly unpredicted manner and thus elicit RPEs. The only RPE could occur at the unpredicted time of the first stimulus that explicitly and quantitatively predicted the reward amounts of the bundle components indicated by the bundle stimulus. Conceivably, in the most simple form, the RPE would reflect the integrated reward amounts of both bundle components relative to the prediction derived from the past trial history. There were three levels of bundle stimulus corresponding to the three IC levels. Thus, appearance of a given bundle stimulus would elicit an RPE relative to the past experienced bundles, weighted by the learning coefficient. Thus, reward prediction errors would have values around −1, 0, and +1 for bundles located on low, intermediate and high ICs, respectively, the variation depending on the learning coefficient. For comparison, the bundle stimulus at each IC level without any RPE would have values of 1, 2 and 3, respectively. Thus, neural responses to the RPE and to the stimulus directly (i. e. without subtraction of prediction) would result in very similar regression slopes (depending on the learning constant used for computing the RPE) and thus be difficult to distinguish from each other. We modelled RPEs with various learning coefficients in the range between 0.1 and 0.9 and for all values found high correlations between RPE and bundle stimulus value at the three IC levels. For example, a learning coefficient of 0.2 in a Rescorla-Wagner model resulted in a Spearman-rank correlation of 0.9337 ± 0.00085 SEM (n=15 bundles x 24 trials = 360 trials x 24 subjects pooled). For this reason, a RPE analysis would not yield new insights and will not be further reported.

## Results

### Implementation of indifference curves

Participants (n = 24) chose between two visual stimuli in repeated trials. Each of the two bundle stimuli represented a two-component bundle that contained the same two milkshakes with independently set amounts (Fig. 1A; see Methods). Thus, we implemented choices between bundles with separate objects (two milkshakes) rather than choices between single objects that each had multiple components. Each stimulus contained two colored vertical rectangles: the blue rectangle represented component A (low-sugar high-fat milkshake); the red rectangle represented component B (high-sugar low-fat milkshake). In each rectangle, a vertically positioned bar indicated the physical amount of each component milkshake, where higher was more.

We examined choices between: (1) a pre-set Reference bundle and (2) a Variable bundle whose component A had a fixed test amount and whose component B varied pseudorandomly. In all 24 participants, choice probabilities followed the component B monotonically. We obtained each indifference point (IP; choice probability *P* = 0.5 for each bundle, indicating equal preference and same utility despite different bundle composition) from a set of six-repetition choices using a probit choice function (Eqs. 1, 1a). We thereby obtained a two-dimensional IP that showed the amounts of the two components of the Variable Bundle between which the participant was indifferent against the constant Reference Bundle. We repeated this procedure, keeping the Reference Bundle constant and increasing the amounts of component A in the Variable Bundle, thus obtaining a set of IPs. All IPs in such a set were equally revealed preferred to, and thus had the same utility as, the constant Reference Bundle.

In each participant, we estimated a total of three sets of IPs (each containing 5 IPs) by pre-setting three different amounts of component B (2 ml, 5 ml or 8 ml with component A always 0 ml) in the Reference Bundle. Each IP defined the trade-off between the two components; it indicated how much of component B the participant was willing give up in order to gain one unit of component A without change of preference. We derived each IC from such a set of five IPs by hyperbolic fitting (Eqs. 2, 2a; Fig. 1B, C). Taken together, the IPs with the continuous ICs represented revealed preferences in a systematic manner, thus implementing the basic concepts underlying this study.

### Behavioral validation of indifference curves

To assess the contribution and validity of IPs (bundles) to the ICs obtained with hyperbolic fits, we performed a leave-one-out analysis. The details of these behavioral analyses were presented before (Pastor-Bernier et al. 2020) and are repeated here for completeness. Briefly, we left out (removed) one IP at a time from the five IPs within one fitted IC (except for the Reference Bundle at x = 0), and then we refitted the IC using the remaining four IPs with the same hyperbolic equation (see Methods, Eqs. 2, 2a). We performed the same kind of leave-one-IP-out analysis separately for each IC in each participant (4 IPs on 3 ICs in 24 participants, resulting in 288 analyses in total).

The refitted ICs resulted in consistent fits in four measures. First, there was no overlap in the refitted IC with any refitted IC at other levels in all 24 participants; thus, the IC levels retained separation despite one IP being left-out. Second, there was no overlap in the 72 refitted ICs with the 95% confidence intervals of other original ICs at different levels; thus, the IC levels retained separation despite one IP being left-out. Third, most refitted ICs (92 %, 66 of 72 ICs) still within the 95% confidence intervals of the original ICs without the eft-out IPs, while the remaining curves (8%, 6 of 72 ICs) showed only some parts of the IC that fell outside the 95% confidence intervals; thus, individual IPs were not overweighted in the ICs. Fourth, the left-out IPs deviated only insignificantly from the refitted ICs (*P* = 0.98 with t-test; *N* = 336; residual: 0.05 ± 0.13 ml in all participants, mean ± standard error, SEM) (Fig. 1D); this result confirmed that individual IPs were not overweighted in the ICs. These four validations demonstrated the robustness and consistency of the hyperbolically fitted ICs in capturing the IPs. Thus, in all participants, the ICs provided valid representations of the three revealed preference levels.

### Neural responses for two-component bundles across and along ICs

During fMRI scanning, the task started with a fixation cross lasting 0.5 s (Fig. 1E). Then, a single two-component visual stimulus appeared in the center of the computer monitor (bundle-on phase); the stimulus predicted delivery of one of the 15 bundles (IPs) composed of two different milkshakes. The physical amount of the milkshakes in the bundle was determined by the participant-specific indifference point (IP) estimated from the binary choice task (see above). The participant received the two bundle milkshakes with the respective amounts indicated by the vertical bars on the stimulus, without choice. That presentation was either followed by a Becker-DeGroot-Marschak (BDM) task within the trial (50% of trials, pseudorandomly selected) or terminated (50% of trials). The BDM bidding served as a mechanism-independent measure of utility estimation, as used before (de Berker et al. 2019; De Martino et al. 2013). In total, each participant performed 360 trials (24 trials for each of the 15 bundles). With the fMRI data we collected, we analyzed the various aspects of neural responses (BOLD signals) to the bundles with several General Linear Models (GLMs) and region-of-interest (ROI) analyses.

We first used GLM1 to identify brain responses that follow the scheme of ICs, namely monotonic increase with higher ICs (or decrease with inverse coding) and insignificant change along the same ICs, as shown in Fig. 2A. Thus, would BOLD signals change monotonically with preference and utility across ICs but vary insignificantly with choice indifference and same utility along ICs? To do so, the individual contrast images (representation of BOLD signal) of each bundle in each participant were grouped according to the IC the bundle belonged to (low, medium, high) and the position of the bundle on each IC (1 - 5, from top left to bottom right).

We used parametric statistical tests (t-test with Flexible Factorial Design) and estimated neuroimages of responses to each of the 15 bundles grouped into the three IC levels or five groups along ICs (see Methods). We found that the striatum, midbrain and OFC showed significantly increasing activation across increasing ICs (high > low IC; map threshold of p < 0.005; t-test) but insignificant variations along individual ICs (exclusive mask map threshold of p < 0.005) (Fig. 2B; Table 1; for effect sizes, see Table 1-1). More specifically, we found small-volume corrected significance in the striatum (peak at [10, 6, −4], z-score = 3.27, 6 mm radius sphere, cluster-level FWE corrected p = 0.041), midbrain (peak at [4, −16, −12], z-score=3.71, 6 mm radius sphere, cluster-level FWE corrected p = 0.048) and OFC (peak at [22, 42, −16], z-score = 3.67, 10 mm radius sphere, cluster-level FWE corrected p = 0.037). (All small-volume corrections in this study were centered on pre-defined coordinates from the Neurosynth meta-analysis database, see Methods). In addition, we found significant activities in other regions, including the insula and cingulate cortex (Table 1). By contrast, we found significant BOLD changes between bundles positioned on same ICs in a number of other, mostly cortical regions (Table 1-2). These changes violated the IC scheme representing the trade-off between the two bundle rewards and were not further explored.

**Table 1.**
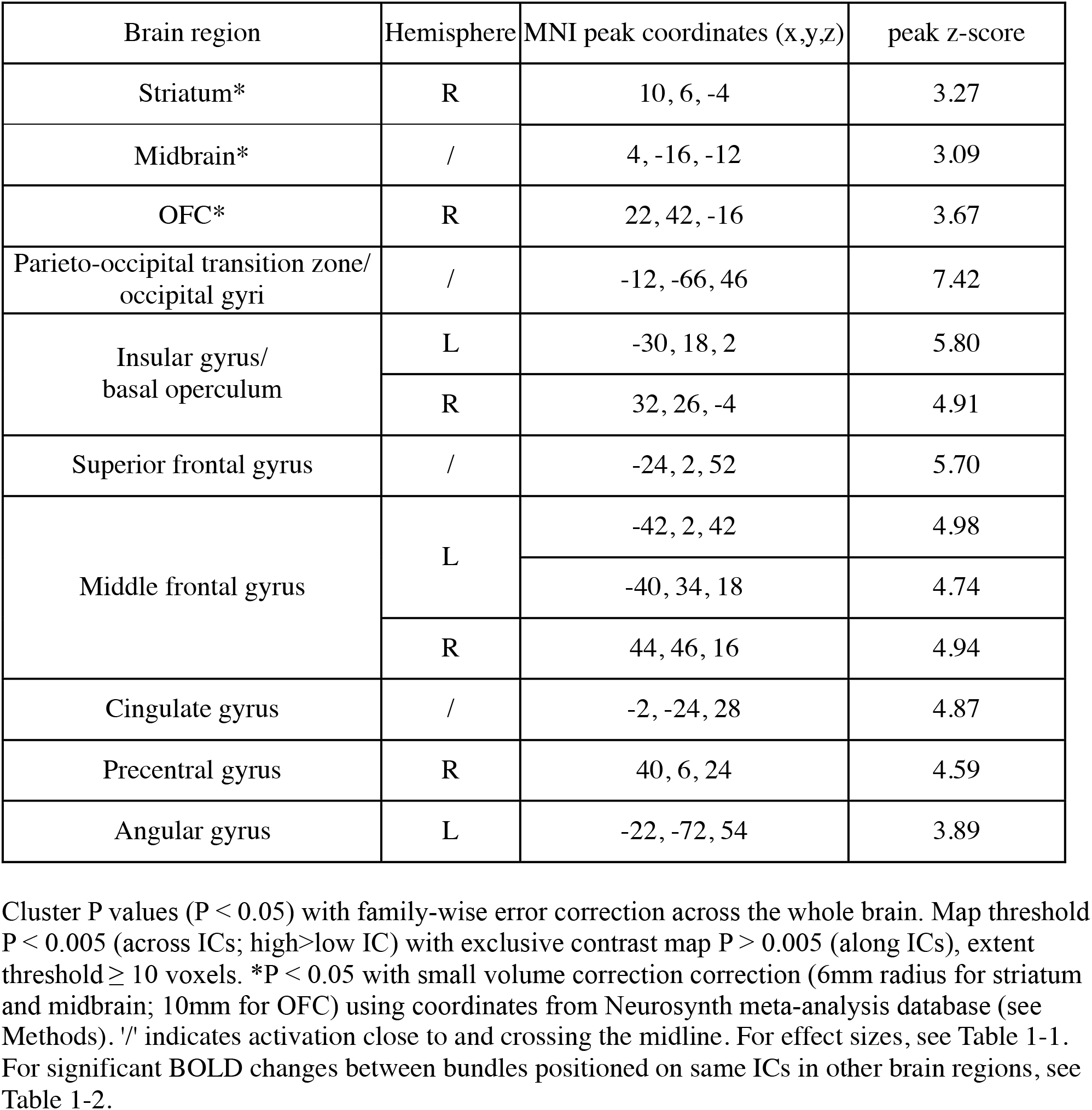
Brain regions activated across but not along indifference curves (ICs) during bundle-on phase (whole-brain analysis with GLM1).

| Brain region | Hemisphere | MNI peak coordinates (x,y,z) | peak z-score |
| --- | --- | --- | --- |
| Striatum* | R | 10, 6, -4 | 3.27 |
| Midbrain* | / | 4, -16, -12 | 3.09 |
| OFC* | R | 22, 42, -16 | 3.67 |
| Parieto-occipital transition zone/<br>occipital gyri | / | -12, -66, 46 | 7.42 |
| Insular gyrus/<br>basal operculum | L | -30, 18, 2 | 5.80 |
|  | R | 32, 26, -4 | 4.91 |
| Superior frontal gyrus | / | -24, 2, 52 | 5.70 |
| Middle frontal gyrus | L | -42, 2, 42 | 4.98 |
|  |  | -40, 34, 18 | 4.74 |
|  | R | 44, 46, 16 | 4.94 |
| Cingulate gyrus | / | -2, -24, 28 | 4.87 |
| Precentral gyrus | R | 40, 6, 24 | 4.59 |
| Angular gyrus | L | -22, -72, 54 | 3.89 |

Cluster P values ( $P < 0.05$ ) with family-wise error correction across the whole brain. Map threshold $P < 0.005$ (across ICs; high>low IC) with exclusive contrast map $P > 0.005$ (along ICs), extent threshold $\geq 10$ voxels. \* $P < 0.05$ with small volume correction (6mm radius for striatum and midbrain; 10mm for OFC) using coordinates from Neurosynth meta-analysis database (see Methods). '/' indicates activation close to and crossing the midline. For effect sizes, see Table 1-1. For significant BOLD changes between bundles positioned on same ICs in other brain regions, see Table 1-2.

**Table 1-1.** Effect sizes for BOLD responses to bundles positioned across and along indifference curves (IC) in striatum, midbrain and OFC (GLM1).

| Brain region | Preference<br>(different utility) | Choice indifference<br>(same utility) |  |
| --- | --- | --- | --- |
|  | Effect across ICs | Effect along ICs<br>(T statistics) | Effect along ICs<br>(F statistics) |
| Striatum | Z = 3.27<br>p = 0.041 | N/A (Fat)<br>N/A (Sugar) | N/A |
| Midbrain | Z = 3.09<br>p = 0.048 | N/A (Fat)<br>N/A (Sugar) | N/A |
| OFC | Z = 3.67<br>p = 0.037 | Z = 2.77, p = 0.188 (Fat)<br>N/A (Sugar) | N/A |

P values refer to small volume corrected BOLD signal (6mm radius for striatum and midbrain; 10mm for OFC) using coordinates from Neurosynth (see Methods). Map threshold $P < 0.005$ , extent threshold $\geq 10$ voxels. Z: peak z-score. Threshold $p = 0.005$ with extent threshold $\geq 10$ voxels, tested with small volume correction. N/A: no cluster of voxels met the statistical criteria.

**Table 1-2.** Brain regions activated along indifference curves (ICs) during bundle-on phase (whole-brain analysis with GLM1 F contrast along ICs).

| Brain region | Hemisphere | MNI peak coordinates (x,y,z) | peak z-score |
| --- | --- | --- | --- |
| Striate area | / | 2, -86, 6 | >8 |
| Inferior frontal gyrus, orbital part/<br>Lateral OFC | R | 46, 50, -2 | 4.63 |
| Inferior frontal gyrus, triangular part | R | 56, 16, 0 | 4.14 |
| Middle temporal gyrus | R | 64, -24, -18 | 4.41 |
| Superior frontal gyrus, medial part | / | 4, 38, 26 | 4.37 |
| Supramarginal gyrus | R | 48, -50, 42 | 4.31 |
| Inferior temporal gyrus | L | -32, -78, -16 | 3.86 |
| Superior temporal gyrus | L | -56, -46, 40 | 3.74 |

Cluster P values ( $P < 0.05$ ) with family-wise error correction across the whole brain. Map threshold $P < 0.005$ (along ICs), extent threshold $\geq 10$ voxels. '/' indicates activation close to and crossing the midline.

To provide further evidence for neural activations following the scheme of ICs, we performed a Spearman rank time course analysis. We first extracted BOLD signals using leave-one-subject-out cross-validated GLM models, which should prevent potential biases with pre-selected peaks (see Methods). Subsequently we used the BOLD signals from peak voxels in each left-out subject to perform Spearman rank analyses. We found that the striatum, midbrain and OFC showed significant Spearman rank correlation coefficients (Spearman’s Rho) between bundles located on different ICs at around 6s after onset of the bundle stimulus (p < 0.05), consistent with the standard time course of haemodynamic response (Fig. 2C). By contrast, only insignificant (p > 0.05) rank coefficients were found at 5 - 7s between bundles located along same ICs in these brain regions, as shown in the sliding-window analysis (Fig. 2D). These time courses followed the revealed preference to bundles across different ICs but failed to differ along the same IC, thus complying with the scheme of ICs that represent revealed preference. Moreover, we extracted beta (slope) coefficients of the BOLD signal at 6s with the ROI coordinates identified by GLM1 and plotted them for three revealed preference levels in bar charts (Fig. 2E). We found a significant difference between high versus low revealed preference level in the midbrain (p = 0.0062), OFC (p = 0.0023), and marginal significant difference in the striatum (p = 0.0533). We also found a significant difference between high versus middle revealed preference level in the OFC (p = 6.8551 x 10^−4^). By contrast, a one-way ANOVA analysis on the beta (slope) coefficients of the BOLD signal indicated insignificant differences between responses to 5 IPs positioned on same ICs in striatum, midbrain and OFC (Fig. 2F). We used F contrasts as the exclusive mask and found small volume corrected significance in striatum (p = 0.041, 6 mm radius sphere) and OFC (p = 0.037, 10 mm radius sphere) but only marginal significance in midbrain (p= 0.051, 6 mm radius sphere) (Fig. 2-1). These activations were also confirmed with the lower threshold of p < 0.001 (Fig. 2-2; T contrast), with small volume corrected significance in striatum (p=0.017, 6 mm radius sphere), OFC (p=0.018, 10 mm radius sphere) and midbrain (p=0.042, 8 mm radius sphere; no significance with 6 mm).

Taken together, these data indicate that activations in several components of the brain’s reward system followed the basic scheme of ICs representing revealed preferences: activation across the ICs but no activation along the same IC.

### Binary comparisons between partial physically non-dominant bundles

According to the concept of ICs, any bundle on a higher IC (farther from the origin) should be preferred to any bundle on a lower IC. Hence, a single-dimensional neural signal reflecting multi-component choice options should vary between any bundle on a higher IC and any bundle on a lower IC. To reflect the proper integration of the two bundle components irrespective of specific physical properties, the neural signal should follow the IC rank even when one component milkshake of the higher-IC bundle is lower than in the lower-IC bundle (partial physical non-dominance). To identify such differences, we used the GLM2. With pairwise comparisons, GLM2 should identify higher responses to revealed preferred bundles with partial physical non-dominance. Thus, GLM2 compared all bundle pairs that fit the following condition within each participant: bundle 1 was located on higher IC but had a lower amount of one component milkshake compared to bundle 2 that was located on a lower IC (Fig. 3A).

The GLM2 analysis demonstrated significant activations in similar regions as with GLM1, where striatum (peak at [16, 6, −6], z-score=3.8, 6 mm radius sphere, cluster-level FWE corrected p = 0.012), midbrain (peak at [4, −16, −12], z-score=2.85, 6 mm radius sphere, cluster-level FWE corrected p = 0.032) and OFC (peak at [24, 42, −16], z-score=3.99, 10 mm radius sphere, cluster-level FWE corrected p = 0.012) showed small-volume corrected significant activations (Fig. 3B). These activations were also confirmed with the lower threshold of p < 0.001 (Fig. 3-1), with small volume corrected significance in striatum (p=0.008, 6 mm radius sphere) and OFC (p=0.004, 10 mm radius sphere). Also, we found significant activities in other regions, including insula, superior frontal gyrus and cingulate, as shown in Table 2.

**Table 2.** Brain regions showing differences (partial physical non-dominance> partial physical dominance) in BOLD signal between partial physically dominating bundles located on different indifference curves (ICs) during bundle-on phase (whole-brain analysis with GLM2).

| Brain region | Hemisphere | MNI peak coordinates (x,y,z) | peak z-score |
| --- | --- | --- | --- |
| Striatum* | R | 16, 6, -6 | 3.8 |
| Midbrain* | / | 4, -16, -12 | 2.85 |
| OFC* | R | 24, 42, -16 | 3.99 |
| Insular gyrus/<br>Basal operculum | L | -30, 24, -2 | 5.55 |
|  | R | 32, 26, -6 | 4.40 |
| Angular gyrus | R | 32, -68, 28 | 5.51 |
| Cerebellum | L | -36, -68, -30 | 4.79 |
| Superior frontal gyrus | / | 22, 2, 54 | 4.75 |
| Occipital gyri | L | -28, -88, 4 | 4.57 |
| Middle frontal gyrus | L | -50, 40, 16 | 4.32 |
|  | R | 46, 42, 14 | 3.76 |
| Inferior frontopolar gyrus | R | 18, 64, -8 | 4.11 |
| Cingulate gyrus | / | -2, -24, 28 | 4.05 |

Cluster P values ( $P < 0.05$ ) with family-wise error correction across the whole brain. Map threshold $P < 0.005$ , extent threshold $\geq 10$ voxels. \* $P < 0.05$ with small volume correction (6mm radius for striatum and midbrain; 10mm for OFC) using coordinates from Neurosynth meta-analysis database (see Methods).

We also performed ROI analyses (coordinates identified by GLM2 with leave-one-subject-out procedure) that calculated betas of partial physical non-dominance (higher revealed preference) and partial physical dominance bundles (lower revealed preference) as described in Methods. For each ROI, we computed three models, which compared bundles pairwise, with low vs. middle, middle vs. high, and low vs. high revealed preference, respectively. Neural beta regression coefficients were extracted at 6 s after the onset of the bundle stimulus, which corresponded to the canonical hemodynamic response. In regard to high vs. low revealed preference level, we found significance in the striatum (p = 0.0459) and OFC (p = 0.0033) when comparing bundles in high IC vs. low IC (Fig. 3C). We also found significance in the striatum (p = 0.0309) and OFC (p = 7.6575 x 10^−5^) when comparing high vs. middle IC bundles. In the midbrain, we found no significance (p > 0.05) in the three comparisons between bundles on low, middle and high ICs (although such a tendency existed in all three comparisons). When plotting Figure 3C using peak voxels from GLM1, we found similar results for all three regions (Fig. 3-2), which is unsurprising as the coordinates were similar between GLM1 and GLM2. Thus, the region-of-interest analysis was robust with GLM1 coordinates for these regions.

Taken together, these pairwise bundle comparisons demonstrated neural coding of partial physical non-dominance bundles as a necessary condition for extracting a scalar neural signal from vectorial, multi-component choice options. These results confirmed compliance with the graphic schemes of ICs demonstrated with GLM1.

### Becker-DeGroot-Marschak (BDM) control of revealed preference

To validate the order of revealed preferences represented by the ICs with an independent estimation mechanism, we used a monetary Becker-DeGroot-Marschak (BDM) bidding task that estimated each participant’s utility for each bundle. In 50% of trials during fMRI scanning, each participant made a monetary BDM bid (UK pence) for one of the 15 bundles, out of a fresh endowment of 20 UK pence in each trial (BDM bidding phase; Fig. 1E). The 15 bundles constituted the indifference points of the ICs that were estimated during the binary choice task with each participant.

The BDM bids followed the order of revealed preference levels across ICs, as demonstrated by significant positive Spearman Rank correlation between the three IC levels and the bid amounts for bundles and confirmed with significant binary Wilcoxon signed-rank tests between the three IC levels (Fig. 4A; blue, green, red). By contrast, there was no correlation between bids for the five bundles and their position along each IC (from top left to bottom right; Spearman Rho = 0.0219; p = 0.6791). Thus, BDM bids increased across the three IC levels but did not change monotonically with bundle position along individual ICs in the population of our participants.

In order to investigate neural mechanisms of BDM bidding and value elicitation, we compared two GLM models: (1) GLM3 to identify brain regions that encoded BDM bids (0 - 20 pence) during the bidding phase, as shown in Fig. 4B; (2) GLM1 to identify brain regions that encoded value elicitation according to IC levels during bundle-on phase, as shown above in Fig. 2B, C, far right.

Analysis with GLM3 demonstrated activation in vmPFC that encoded BDM bids during the bidding phase (Fig. 4B left; peak at [6, 44, 0], z-score = 4.10, whole-brain corrected with cluster-level FWE corrected p = 0.002), together with other brain regions (Table 3). Further ROI analysis showed significant rank correlation between vmPFC activation and BDM bids at around 6 s after BDM cue onset (Fig. 4B right; bidding phase; p < 0.05; Spearman’s Rho), consistent with the expected haemodynamic response function. By contrast, analysis with GLM1 showed significant, small-volume corrected activation in OFC that indicated its involvement in encoding IC levels during the bundle-on phase (Fig. 2B; far right). The ROI analysis showed significant rank correlation between OFC activation and IC levels at around 6 s after bundle onset (Fig. 2C; far right; bundle-on phase; p < 0.05; Spearman’s Rho). In addition, with GLM3, we found small-volume corrected significant activation in the dorsal striatum (Fig. 4-1; peak at [12, 12, 0], z-score = 3.53, 6 mm radius sphere, cluster-level FWE corrected p = 0.008) encoding the BDM bids but not in the ventral striatum (p > 0.1).

**Table 3.** Brain regions with BOLD responses correlating with BDM bids during the bidding phase (whole-brain analysis with GLM3).

| Brain region | Hemisphere | MNI peak coordinates (x,y,z) | peak z-score |
| --- | --- | --- | --- |
| vmPFC | / | 6, 44, 0 | 4.10 |
| Dorsal striatum* | R | 12, 12, 0 | 3.53 |
| Insular gyrus/<br>basal operculum | R | 30, 24, -2 | 5.21 |
| Inferior frontal gyrus, opercular<br>part | R | 46, 8, 22 | 4.76 |
| Occipital gyri | L | -36, -90, -2 | 4.67 |
| Superior parietal lobule | L | -30, -56, 46 | 4.14 |
| Superior frontal gyrus | / | 2, 26, 42 | 4.05 |
| Middle frontal gyrus | R | 38, 36, 18 | 3.93 |
| Postcentral gyrus | R | 32, -36, 48 | 3.62 |
| Inferior frontopolar gyrus | R | 22, 56, -4 | 3.53 |

Cluster P values ( $P < 0.05$ ) with family-wise error correction across the whole brain. Map threshold $P < 0.005$ , extent threshold $\geq 10$ voxels. \* $P < 0.05$ with small volume correction (6 mm radius) using coordinates from a previous study with BDM bidding (De Martino et al., 2009; see Methods).

Taken together, BDM bidding provided a good validation of the estimated levels of revealed preference represented by ICs. However, and interestingly, revealed preference levels and BDM bids were encoded in different regions of the frontal cortex and striatum.

## Discussion

We systematically tested characteristics of scalar neural responses to vectorial, multi-component bundles. We estimated indifference points (IPs) by asking human participants to choose between two bundles. Each bundle contained the same two separate objects (milkshakes) rather than consisting of single objects that each had multiple components. Our behavioral results (Pastor-Bernier et al. 2020) showed that preference relationships among multi-component choices were reliably represented by systematic ICs, as a prerequisite for testing the underlying neural mechanisms. In fMRI scans with GLM and post-hoc ROI analyses, we identified brain regions whose activations correlated with levels of revealed preference. The GLM1 and post-hoc Spearman rank analysis demonstrated activations in the ventral striatum, midbrain and OFC that reflected revealed preference levels across ICs (changing utility) but failed to vary along equal-preference ICs (same utility despite different bundle composition). The GLM2 specifically dissociated revealed preference from physical dominance and showed consistent results with those from GLM1. A mechanism-independent control with a Becker-DeGroot-Marschak (BDM) bidding task confirmed the validity of ICs for representing revealed preference levels. Interestingly, however, BDM bidding was associated with activations in vmPFC and dorsal striatum rather than the previously identified reward structures following IC levels. Together, these data demonstrate systematic, single-dimensional neural activations in the striatum, midbrain and OFC that reflect preferences for, and utility of, vectorial multi-component choice options.

Scalar neural activations from vectorial choice options are only the most simple way to represent value integrated from multiple components. Other plausible but less straightforward ways might be ensemble coding composed of multiple heterogeneous signals representing only single components of multi-component options, as seen in individual OFC neurons (Pastor-Bernier et al. 2019). Future neuroimaging studies may address such issues.

In our binary choice task, we elicited revealed preferences with repeated, psychophysically controlled choices (Pastor-Bernier et al. 2020; Green, & Swets 1966). Such a multi-trial, stochastic approach is well conceptualized (McFadden & Richter 1990; McFadden 2004), fulfils statistical requirements of neural research, corresponds to standard choice functions (Sutton, & Barto 1998), and allows comparison with animal neurophysiology (Pastor-Bernier et al., 2017). These methods delivered varying choice probabilities (stochastic choices) instead of single selections (deterministic choices).

Economic choice experiments often involve substantial but imaginary sizes or amounts of consumer items and money, or use random singular payouts (Simonson, 1989; Tversky & Simonson 1993; Rieskamp el al., 2006). By contrast, our payout schedule fit the requirements of neuroimaging and involved tangible and consumable rewards over hundreds of trials, while also controlling for satiety. The behavioral choices resembled small daily activities, such as drink and snack consumption. In this way, we obtained three well-ordered ICs for each participant that provided accurate and systematic representations of preferences for multi-component bundles, without involving imagined items or monetary reward (Pastor-Bernier et al. 2020).

We used the BDM task as an authoritative, mechanism-independent control for eliciting subjective values, thereby providing an additional validating mechanism for the revealed preferences elicited in our binary choice test. The value estimating mechanism for BDM bids differs substantially from the one for revealed preference ICs. The truthful revelations (incentive compatibility) of BDM makes this mechanism an essential tool in experimental economics that is becoming more popular in human decision research (Plassmann et al., 2007; Medic et al., 2014; Zangemeister et al., 2016). The elicited BDM bids correlated well with the revealed preference levels (Pastor-Bernier et al., 2020) and thereby validated in a mechanism-independent manner the empirically estimated IPs used during fMRI (in which the participants performed the BDM task). Previous neuroimaging studies showed activations in ventromedial prefrontal cortex that correlated with BDM bids (Chib et al., 2009; McNamee et al., 2013). Our experimental design dissociated value elicitation by bundles and by BDM bidding. We confirmed the BDM activations in vmPFC and found that the two mechanistically different tasks activated different regions in both prefrontal cortex and striatum; responses to the bundles followed the IC scheme (different activations across but not within ICs) in OFC and ventral striatum, whereas BDM bidding activated vmPFC and dorsal striatum. Previous studies showed that vmPFC activity can reflect value derived from both rating measures and can distinguish between preferred and non-preferred options irrespective of task demands (Lebreton et al., 2009; Lopez-Persem et al., 2020). Thus, the conditions under which vmPFC encodes value, and the precise form of value-elicitation that best explains vmPFC activity are valuable topic for future studies..

Previous studies tested neural mechanisms of human choice of bundles with multiple components, such as payoff amount and probability (Chau et al. 2014), quality and quantity of goods (de Berker et al. 2019), money and time (Gluth et al. 2017), and food components (Suzuki et al. 2017). Nevertheless, none of these studies tested bundles that were positioned along modelled ICs (i.e. eliciting choice indifference) and thus failed to test the crucial trade-off that demonstrates the graded and well-ordered manner of single-dimensional preferences for multi-dimensional choice options. Without this information, we would not know how a scalar neural response may arise from graded changes of vectorial, multi-component bundles. Our study, testing 5 bundles on each IC, addressed this problem and identified the brain regions that showed this kind of neural response.

Although we tested the emergence of single-dimensional neural signals for multi-dimensional bundles in a systematic and concept-driven way, there were limitations with our experimental design. First, both bundle components had the same type of primary reward (milkshakes). It would be interesting to study whether the same brain regions would encode different types of rewards and follow the formalisms of ICs, including the graded trade-off. For instance, future research may compare monetary rewards with primary nutrient rewards. Second, we only demonstrated neural responses with the typical convex ICs. It would be interesting to study whether different brain regions might encode preferences with different shapes of ICs. Such work may test participants’ choices with linear or concave ICs. Third, we did not test the influences of prior experience on current decisions. Previous studies (Schultz 1998; van den Bos et al. 2013; Lopez-Persem et al. 2016) showed that choices could be influenced by previous experience and be updated by reinforcement learning. Future research may include multi-component choice options during fMRI scanning to study multi-component reinforcement learning. Lastly, we only demonstrated fMRI BOLD responses, and future neurophysiology research should confirm the coding of revealed preference at a single neuron level in human patients with intracerebrally implanted electrodes, similar to our recently investigated neuronal encoding of revealed preference in monkey orbitofrontal cortex (Pastor-Bernier et al. 2019). To conclude, while we showed brain activation with bundles in a formal but standard revealed preference setting (convex ICs, primary reward), it is desirable to know how human brains encode revealed preference in a larger variety of situations.

The reward circuit including the striatum and midbrain is known to participate in reward anticipation and learning, including reward prediction error (Diederen et al. 2017). In monkeys, midbrain dopamine neurons encode values for predicted rewards in economic decision tasks (Lak et al. 2016; Schultz et al. 2017). Similar to the midbrain and striatum, previous work showed the involvement of the human mid-OFC in valuation of primary nutrient reward (Grabenhorst et al., 2010) and monetary reward (Kahnt et al. 2014). Remarkably, the neural activity in OFC elicited here, in response to visual cues predicting liquid rewards with varying sugar and fat components, closely matched the coordinates observed previously (Grabenhorst et al., 2010) in a study in which subjects orally sampled very similar liquid rewards. Thus, this area of OFC seems to be involved both in reward valuation during oral consumption of primary nutrient rewards and in the economic valuation of visually cued choice options. In non-human primates, OFC neurons encode reward prediction (Tremblay & Schultz 1999; Padoa-Schioppa & Assad 2006) and follow revealed preferences for multi-component bundles (Pastor-Bernier et al. 2019). In the current study, we used a concept-driven design and found that neural responses in the striatum, midbrain and OFC integrated multiple bundle components in a way that followed the ICs scheme (changing across ICs but being similar along equal-preference ICs). Moreover, we demonstrate the involvement of the midbrain in multi-component decision making for the first time. Overall, our results show the involvement of principal reward structures of the brain in integrating the multiple components of vectorial bundles into single-dimensional neural signals that are suitable for economic decision making.

Besides the primary reward circuit (midbrain dopamine neurons, OFC, striatum, amygdala), other brain regions are also involved in economic decision making. Previous studies in multi-component decision making suggested the involvement of the cingulate, prefrontal cortex and insula in value elicitation (Kurtz-David et al. 2019; Busemeyer et al. 2019). Consistent with these studies, we also found significant activation in these regions. As shown in Table 1 and Table 2, the BOLD signals identified by GLM1 and GLM2 showed that these regions also encode bundle values during the bundle-on phase, together with the striatum, midbrain, and mid-OFC. Our results are consistent with these previous studies, suggesting that a considerable number of brain regions also play a role in multi-component decision making.

## Conflict of interest

The authors declare no conflicts of interest.

## Acknowledgements

We thank Charles R. Plott for discussions and conceptual support, Steve Edgley for help and logistic support, Simone Ferrari-Toniolo for comments on experimental economics, Jae-Chang Kim and Putu Agus Khorisantono for suggestions on fMRI analysis, and Arkadiusz Stasiak for computer support. The Wellcome Trust supported this work (WT 095495, WT 204811).

## Notes

### Competing Interest Statement

The authors have declared no competing interest.

### Summary of Updates

Several small corrections

